# A systems approach to refine disease taxonomy by integrating phenotypic and molecular networks

**DOI:** 10.1101/219089

**Authors:** Xuezhong Zhou, Lei Lei, Jun Liu, Arda Halu, Yingying Zhang, Bing Li, Zhili Guo, Guangming Liu, Changkai Sun, Joseph Loscalzo, Amitabh Sharma, Zhong Wang

**Affiliations:** School of Computer and Information Technology, Beijing Jiaotong University, Beijing 100044, China.; Institute of Information on Traditional Chinese Medicine, China Academy of Chinese Medical Sciences, Beijing, 100700, China.; Institute of Basic Research in Clinical Medicine, China Academy of Chinese Medical Sciences, Beijing, 100700, China.; Channing Division of Network Medicine, Department of Medicine, Brigham and Women’s Hospital, Harvard Medical School, 181 Longwood Avenue, Boston, Massachusetts 02115, USA.; Department of Medicine, Brigham and Women’s Hospital, Harvard Medical School, 75 Francis Street, Boston, MA 02115, USA.; Jiaxing Traditional Chinese Medicine Affiliated Hospital of Zhejiang Chinese Medical University, Jiaxing 314000, China.; Department of Biomedical Engineering, Dalian University of Technology, Dalian, 116024, China

**Author notes:** These authors contributed equally to this work (Xuezhong Zhou, Lei Lei, Jun Liu, Arda Halu).

## Abstract

The International Classification of Diseases (ICD) relies on clinical features and lags behind the current understanding of the molecular specificity of disease pathobiology, necessitating approaches that incorporate growing biomedical data for classifying diseases to meet the needs of precision medicine. Our analysis revealed that the heterogeneous molecular diversity of disease chapters and the blurred boundary between disease categories in ICD should be further investigated. Here, we propose a new classification of diseases (NCD) by developing an algorithm that predicts the additional categories of a disease by integrating multiple networks consisting of disease phenotypes and their molecular profiles. With statistical validations from phenotype-genotype associations and interactome networks, we demonstrate that NCD improves disease specificity owing to its overlapping categories and polyhierarchical structure. Furthermore, NCD captures the molecular diversity of diseases and defines clearer boundaries in terms of both phenotypic similarity and molecular associations, establishing a rational strategy to reform disease taxonomy.

## Introduction

Disease taxonomy plays an important role in defining the diagnosis, treatment, and mechanisms of human diseases. The principle of the current clinical disease taxonomies, in particular the International Classification of Diseases (ICD), goes back to the work of William Farr in the nineteenth century and is primarily derived from the differentiation of clinical features (e.g. symptoms and micro-examination of diseased tissues and cells)(Council et al. 2011). Despite its extensive clinical use, this classification system lacks the depth required for precision medicine with the limitations of its rigid hierarchical structure and, moreover, it does not exploit the rapidly expanding molecular insights of disease phenotypes. For example, many diseases (e.g. cancer, chronic inflammatory diseases) in the current disease taxonomies have either high genetic heterogeneity (Bianchini et al. 2016; McClellan and King 2010) or manifestation diversity(Arostegui et al. 2014; Jeste and Geschwind 2014; Mannino 2002), which give little basis for tailoring treatment to a patient’s pathophysiology. Furthermore, disease comorbidities (Hu, Thomas, and Brunak 2016; Lee et al. 2008; Hidalgo et al. 2009), temporal disease trajectories(Jensen et al. 2014) in clinical populations, various molecular relationships between disease-associated cellular components and their connections in the interactome (Blair et al. 2013; Goh et al. 2007; Barabasi, Gulbahce, and Loscalzo 2011; Rzhetsky et al. 2007; Zhou et al. 2014), and many successful drug repurposing cases (Li and Jones 2012; Chong and Sullivan 2007; Ashburn and Thor 2004; Wu et al. 2016; Evans et al. 2005) altogether demonstrate the vague boundary between different diseases in current disease taxonomies. Moreover, the deep understanding of diseases based on the advances in disease biology, bioinformatics, and multi-omics data necessitates the reclassification of disease taxonomy (Mirnezami, Nicholson, and Darzi 2012). In the past decade, efforts to reclassify diseases based on molecular insights have increased with studies related to molecular-based disease subtyping in different disease conditions, such as acute leukemias(Golub et al. 1999; Alizadeh et al. 2000), colorectal cancer(Dienstmann et al. 2017), oesophageal carcinoma(Cancer Genome Atlas Research et al. 2017), pancreatic cancer(Bailey et al. 2016), cancer metastasis(Chuang et al. 2007), neurodegenerative disorders(Mann et al. 2000), autoimmunity disorders(Ahmad, Marshall, and Jewell 2003), multiple cancer types across tissues of origin(Hoadley et al. 2014), and a network-based stratification method for cancer subtyping(Hofree et al. 2013). Further insights will arise from integrating all types of biomedical data with a single framework to exploit disease-disease relationships. Data integration methods that utilize multiple types of data, including ontological and omics data, have been used to classify and refine disease relationships (Gligorijevic and Przulj 2015; Menche et al. 2015; Gligorijevic, Malod-Dognin, and Przulj 2016). Despite these efforts, the development of a molecular-based disease taxonomy that links molecular networks and pathophenotypes still remains challenging (Menche et al. 2015; Hofmann-Apitius et al. 2015; Jameson and Longo 2015). Here, we aim to refine a widely used clinical disease classification scheme, the ICD. To achieve this, we first quantify the Category Similarity (CS) among the ICD chapters using ontology-based similarity measures and investigate the molecular connections of disease pairs in the same ICD chapters. Furthermore, we seek the correlation between category and molecular similarity, and check for the heterogeneity of molecular specificity and correlated boundary between categories in ICD taxonomy. Finally, we construct a new classification of diseases (NCD) with overlapping structures. The aim is to provide clear boundaries between distinct diseases belonging to different categories using a new disease classification scheme (Fig.1 & Fig. S3).

**Figure 1.**
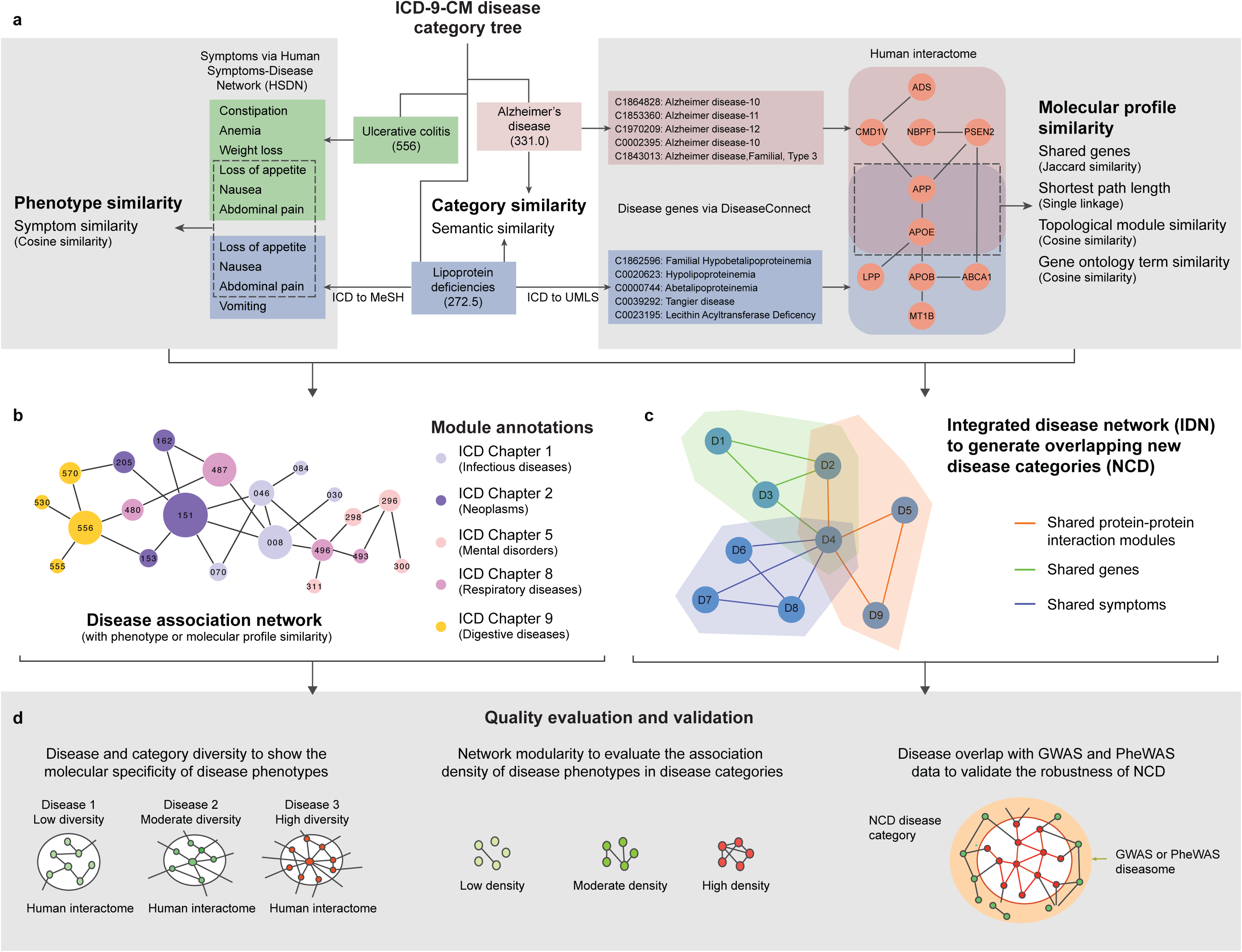
Overview of the new disease taxonomy construction and validation. **a.** Similarity calculation between the disease pairs in ICD taxonomy, including the calculation of 1) category similarity; 2) Phenotype similarity (based on ICD-MeSH term mapping) and 3) Molecular profile similarities (based on ICD-UMLS term mapping) of disease pairs in ICD; **b.** Module or community annotations of disease association network by chapters in ICD or NCD. We generate disease association network, in which nodes represent diseases and the link weights represent their corresponding phenotype or molecule profile similarities. The module annotations of the disease network correspond to ICD chapters or NCD categories; **c.** Construction of integrated disease network (IDN) and generation of NCD. The links of IDN are fused from the multiple similarities (e.g. phenotype similarity and shared gene similarity). Based on IDN, NCD is generated by community detection algorithms with overlapping disease members; **d.** Quality evaluation and validation of ICD and NCD. The molecular specificity (or inverse molecular diversity) and network modularity are used for evaluation and comparison of the quality of two disease taxonomies. Furthermore, we validate the robustness of NCD with two independent phenotype-genotype association datasets, namely GWAS and PheWAS.

## Results

### Category similarity of ICD taxonomy

We curated 1,883 distinct ICD disease codes (Table S1) from the 5-level tree structure of 14,292 ICD-9-CM codes, as well as high confidence protein-protein interactions consisting of 15,551 nodes and 218,409 edges (Franceschini et al. 2013). We compiled 153,277 distinct disease-gene associations between 4552 distinct diseases in UMLS codes and 14975 genes reported in the DiseaseConnect database (Liu et al. 2014) (Fig. S3). Next, by manually mapping the DiseaseConnect identifiers to ICD codes, we obtained 160,754 disease-gene records involving 1,883 distinct ICD codes and 14,906 genes (Fig. S1-2 and Data S1).

To evaluate the closeness of two diseases in the ICD tree structure, we applied an established semantic similarity algorithm (called Category Similarity, or CS). The CS measure, which is a similarity measure based on the information content (IC) signifying how specific a term is, applies to any categorization scheme that has a rooted tree structure, including the ICD-9-CM disease classification. Information theoretic measures such as IC have been used in the context of ICD-9-CM previously(Dahlem, Maniloff, and Ratti 2015). The CS measure takes as input two concepts c1 and c2 and outputs a numeric measure of similarity. If two ICD codes are very close (as in having a very specific common parent code) in the taxonomic tree structure, then the CS would be ~ 1 (Methods, Supplementary Materials (SM) section 2.1). We obtained a disease network comprising 1,883 nodes (representing ICD codes) and 154,563 links, where the edge weight reflects the CS values. The higher the CS, the greater the similarity between diseases whose code positions are adjacent in the ICD tree. The CS distribution showed that most disease pairs (135,271, 87.52%) had category similarities between 0.2-0.5 (Fig. S4a). Disease pairs within this CS range mostly belong to different disease subcategories in the same chapter (e.g. diseases of other endocrine glands and disorders of thyroid gland). For example, the disease pair: type 2 diabetes (ICD: 250.00) and simple goiter (ICD: 240.0), which is in ICD chapter 3, has CS 0.37. However, there do exist disease pairs with high category similarities, such as type 2 diabetes (ICD: 250.00) and type 1 diabetes (ICD: 250.01) with CS 0.83.

While the ICD classification was derived from clinical observations and does not necessarily reflect the connections among the molecular components of diseases, it is informative to quantify to what extent it carries molecular information. To address this, we investigated the correlations of CS of disease pairs with 1) the degree of shared genes and shared clinical phenotypes, 2) GO term (Cell Component, Molecular Function, Biology Process) similarity(Mistry and Pavlidis 2008), and 3) topological similarity (i.e., minimum shortest path length and molecular module similarity) among them (Methods, SM section 2.2). We found that close disease codes (disease pairs with a high CS) actually have higher clinical phenotype similarity (Methods, SM section 2.3), which adheres to the construction principle of ICD taxonomy based on symptom phenotypes (Fig. S4b, PCC=0.960, 95% CI=[0.854, 1.000], p=2.079e-05). Furthermore, we observed strong correlations between higher CS bins compared to lower CS bins for molecular profiles (Fig. S4c-i and Table S2. See Methods, SM section 2 for detailed information). In particular, we observed that in addition to the strongly positive correlations, the percentage overlap of disease pairs with shared genes was generally larger than the random controls (Fig. S4c and S4d, see Methods, SM section 2.4), particularly in the CS region of [0.6-0.9]. The top 10 disease pairs with the largest number of shared genes are all from Chapter 2, which consists of cancer types. This might reflect the fact that cancers are the most studied and complex disease phenotypes involving various gene mutations (Table S3, see Methods, SM section 2 for detailed information). Overall, these findings indicate that diseases in the same ICD chapter tend to have a higher degree of shared genes, and the closer their positions in the ICD tree, the higher is the degree of shared genes.

### Heterogeneity of molecular specificity in ICD taxonomy

We had previously proposed that diseases with diverse clinical manifestations also have more diverse underlying cellular networks (Zhou et al. 2014). Thus, we measured the maximum betweenness of disease-related genes in the protein-protein interaction (PPI) network to quantify the molecular diversity (the inverse of specificity) of each disease, as described previously (Zhou et al. 2014) (see Methods, SM section 3), where a high maximum betweenness indicates a high molecular diversity (MD). While both node diversity (Zhou et al. 2014) and betweenness centrality can be used to determine disease diversity, we choose to use the more established and well-known of these measures, i.e. betweenness centrality. For example, the MD of Alzheimer’s disease could be represented by the maximum betweenness of its related genes (i.e., the betweenness of the APP gene) in the PPI network (Fig. S5a). We observed that the MD of diseases in the ICD taxonomy is heterogeneous, with MD values varying from 10^-8^ to 10^-2^ with a median value of 8.93e-04 (Fig. 2a and Data S3). The top two disease chapters with the highest median MD were Chapter 2 (3.87e-03) and Chapter 1 (1.31e-03) (Fig. 2b). Furthermore, we found that neoplasm (Chapter 2) and infectious disease (Chapter 1) categories tended to have higher MD compared to their complementary categories (Neoplasms vs. Non-Neoplasms p<2.2e-16, Infectious diseases vs. non-infectious diseases p=2.0e-02, Fig. S5b-c) and random controls. We also found that disease categories with unspecified conditions had higher MD compared to disease categories with specific conditions (p=9.75e-03, Fig. S5d, Data S4) and its random control. These results indicate that the molecular diversity of the diseases in neoplasms, infectious diseases, and “Other/unspecified diseases” categories is still an elusive issue that should be addressed. A detailed discussion of disease cases is offered in SM section 3, Data S5 & Table S4-S5.

**Figure 2.**
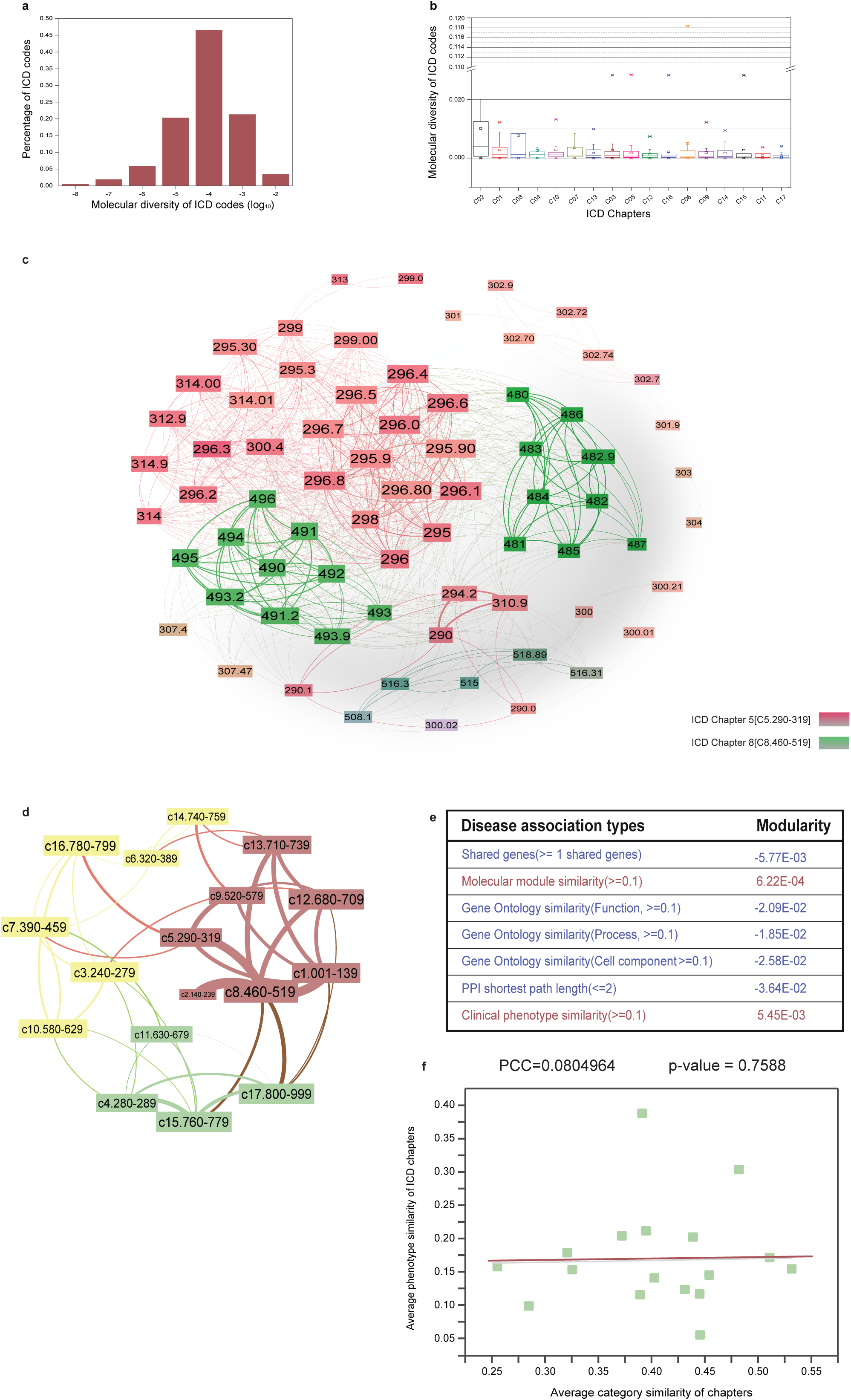
Lack of molecular specificity in ICD taxonomy and the blurred boundary between disease categories in ICD taxonomy. **a.** The distribution of molecular diversity of 1,883 ICD diseases; **b.** The boxplot of molecular diversity of 17 ICD chapters (ordered by median values); **c.**The disease network with shared genes in which the diseases belong to Chapter 5 and Chapter 8. The ICD codes 295, 296, in Chapter 5 have dense relationships to the ICD codes in Chapter 8; **d.** The disease category network with shared genes. The nodes indicate the disease chapters and the weights of edges represent the edge densities between disease chapter pairs; the nodes with same color are considered as a chapter cluster, which is detected by community detection algorithm; **e.** Modularity of disease networks with chapter as module annotations; **f.** The correlation between category similarity and phenotype similarity of ICD chapters.

### The blurred boundary between ICD categories

In the current ICD taxonomy, we observed many instances where there exists a significant number of links between diseases in different chapters, comparable to the number of links between diseases within the same chapter (Table S6 & Fig. 2d, see Methods & SM section 4). For example, strong shared-gene relationships were detected between respiratory diseases (Chapter 8) and mental, behavioral, and neurodevelopmental disorders (Chapter 5) (Fig. 2c-d, more examples shown in SM section 4, Table S7-9). In addition, by calculating the shared molecular connections between diseases in the context of chapters, we could detect 768 diseases with a significant number of shared genes with diseases other than those in their own chapters (Data S6 & SM section 4). To further quantify the molecular boundaries between the disease categories in ICD disease taxonomy, we evaluated the modularity, a structural measure of the tendency of the network to form close-knit communities (see Methods, SM section 2.5) generated by either shared molecular profiles or shared phenotypes. When we mapped ICD chapters as grouping annotations on the various disease networks (filtered by with appropriate weight thresholds) and calculated the modularity, we obtained very low modularity values (Fig. 2e). Since modularity is a widely used measure to validate the quality of partitions/module structures in complex networks, this means that the grouping of ICD chapters does not agree with the natural topological groupings of their corresponding molecular networks (disease modules). This finding gives strong evidence for the blurred disease boundaries of the ICD taxonomy. Furthermore, although the modularity of disease networks with shared phenotypes (similarity>=0.1) is slightly positive, the weak correlation (PCC=0.08, p-value=0.7588) between phenotypic similarity and CS of disease pairs in each chapter (Fig. 2f) indicates that ICD taxonomy does not adequately incorporate phenotype similarity knowledge into disease category structures. Thus, these observations indicate that the strict tree structures in which terms can only have one lineage (Cimino 2011) in the conventional ICD taxonomy may be inefficient, given contemporary knowledge of disease pathobiology, and, therefore, should be refined to be polyhierarchical in structure.

### Polyhierarchical mapping of diseases using molecular module similarity

It has been proposed that if two disease modules overlap in the molecular interaction network, local perturbations in one disease might disrupt the biological pathways in the other disease, which results in shared pathobiological characteristics (Menche et al. 2015). We observed a strong positive correlation between CS and molecular module similarity (MS) (see Methods, SM section 5.1) of diseases, which indicates that two diseases with higher MS would be more closely localized in the disease category (Fig. 3a, PCC=0.887, 95% CI=[0.584,0.973], p=6.12e-04; 3b, PCC=0.974, 95% CI=[0.889,0.994], p=2.08e-06). Here, we investigated the feasibility of utilizing the MS between disease pairs to predict the multiple categories for diseases. Using heuristic rules incorporating the positive correlation between CS and MS, we could predict the closeness of the category location (e.g., shared root parent code) of each given disease pair with positive MS (see Methods, SM section 5.1). In particular, using the 598,420 disease pairs with positive MS values (Data S7), we generated 2,057 predicted additional category results for 722 out of 1,883 disease codes (38.3%) in which each disease code had ~4 categories on average (Data S7&8). We found that the number of predicted categories positively correlated with the MD of the original disease codes (Fig. 3c, PCC=0.547, 95% CI=[0.514, 0.578], p<4.94e-324; External validations see SM 5.2), which indicates that diseases with multiple pathogenic pathways could be captured by polyhierarchical mapping. For example, the 20 diseases in Chapter 8 (i.e. Diseases of the Respiratory System) have been predicted to belong to over five additional chapters, such as neoplasms, infectious diseases, and diseases of the skin and subcutaneous tissue (Fig. 3d), which is consistent with the heterogeneous pathogenesis of COPD and asthma (Grainge et al. 2016; Sharma et al. 2015). A detailed discussion on the polyhierarchial map of the mental disorders is offered in SM section 5.3 (Fig. S8). Furthermore, we found that the predicted category framework had higher phenotype similarity than diseases with shared root codes in the original ICD chapters (see SM section 5.2, median: 0.0703 vs. 0.0563; mean: 0.125 vs 0.109; p <2.2e-16, Fig. 3e. This observation helps to establish that the predicted category results are of higher quality than ICD with respect to their phenotype homogeneity.

**Figure 3.**
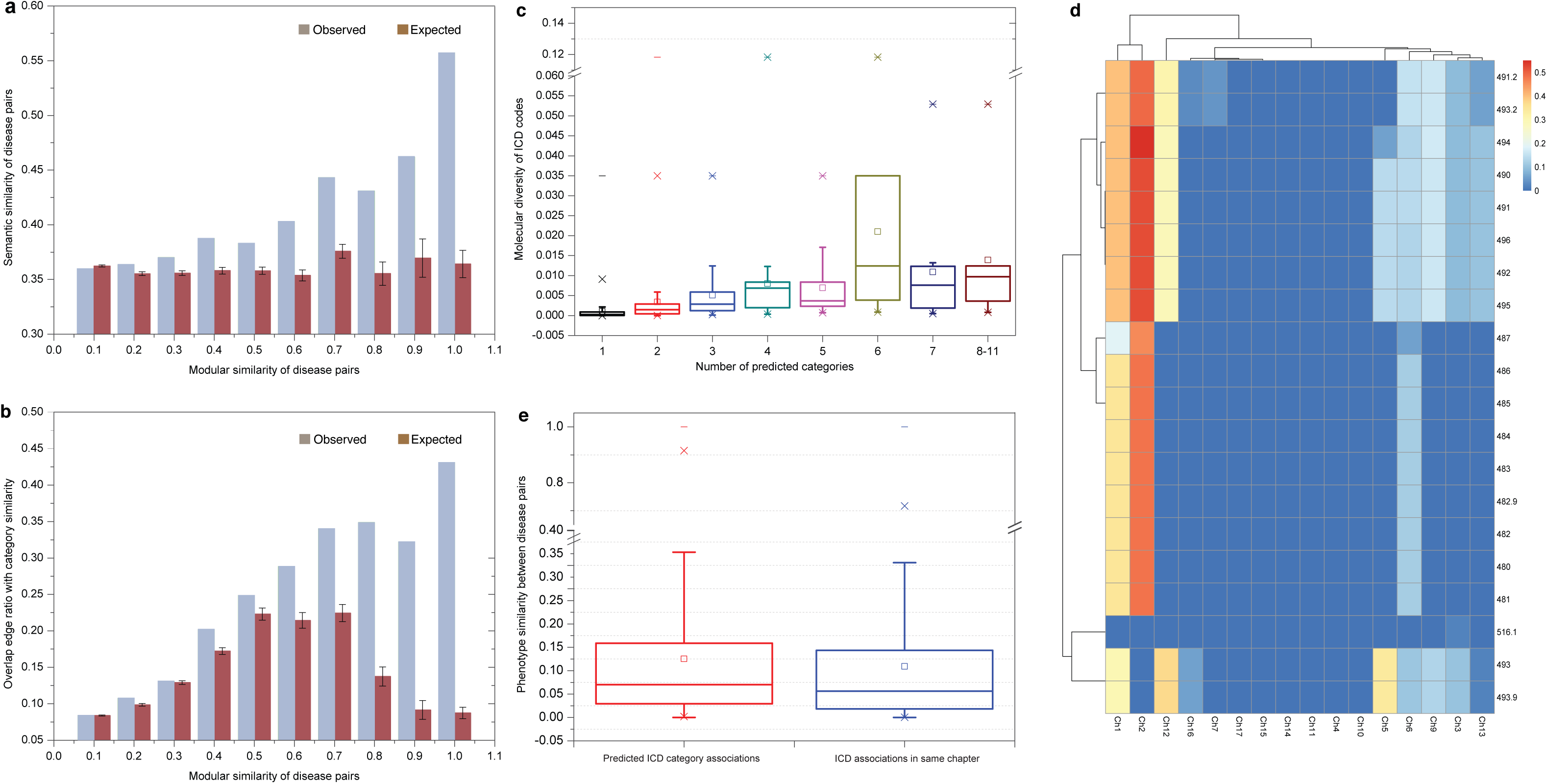
Polyhierarchical map prediction of ICD taxonomy based on molecular module similarity. **a.** Correlation between mean semantic (category) similarity and mean modular similarity of disease pairs; **b.** Correlation between overlapping edge ratio with category similarity and modular similarity of disease pairs; **c.** Correlation between predicted category number and molecular diversity of ICD codes in Chapter 14 (PCC: 0.438; 95% CI: [0.401, 0.474]; p<2.2e-16); **d.** Polyhierarchical map of the disease codes in Chapter 8, indicating that the 20 disease codes in Chapter 8 have significant associations with two disease category clusters: 1) Chapter 1 (infectious disease) and Chapter 2 (neoplasms); 2) Chapter 3 (endocrine, nutritional and metabolic diseases and immunity disorders), Chapter 5 (mental disorders), Chapter 6 (nervous diseases), Chapter 9 (digestive diseases), Chapter 12 (skin and subcutaneous disease) and Chapter 13 (musculoskeletal system and connective tissue diseases); **e.** The boxplots of phenotype similarity of predicted disease pairs and original ICD disease pairs in the same top-level chapters (p<2.2e-16, Wilcoxon test).

### Integrated disease network for overlapping disease classification

To extend and redefine disease concepts by discovering additional categories of a disease, and thereby generate a novel disease taxonomy, we next constructed an integrated disease network (IDN) with:

(a) Shared clinical phenotypes

(I) shared symptoms

and

(b) Molecular profiles

(I) shared genes and molecular module similarity
(II) shortest path lengths in the PPI network

based on a systematic integration process to filter out possible false positive associations (see Methods, SM section 6, Fig. S9 and Fig. S11a), which includes 1,857 diseases and 35,114 links (Data S9).

We then applied high performance community detection algorithms to identify overlapping community structures in the IDN (see Methods, SM section 7 and Fig. S11a). In particular, we first used BigClam (see Methods) since this method is able to detect overlapping communities whereby a disease can belong to multiple communities, in line with our main premise of creating a molecular based flexible disease classification. This resulted in 223 disease sub-categories with overlapping diseases as members (Fig. S11a and Data S10), which included 1,797 distinct diseases from the ICD taxonomy. These 223 disease sub-categories contain different numbers of ICD codes, ranging from 5 to 168 (Fig. S12), therefore, they represent different levels of disease categories similar to ICD chapters and their sub-categories. Next, to develop a more unified view of the disease category quality, we used the well-established BGLL method, which detects non-overlapping communities, to cluster these 223 sub-categories further into 17 non-overlapping, distinct parts, such that these represent the 17 new chapter-level categories (called new chapters, or NCs) using the shared ICD codes (see Methods, SM section 7.2, Fig. S11b). Overall, this clustering order effectively ensures distinct top-level categories that have overlapping subcategories. The resulting 17 NCs contain different numbers of sub-categories ranging from 4 to 25, or of diseases ranging from 53 to 369 (Fig. S11c). We denote the 17 NCs together with their 223 disease sub-categories as our new overlapping disease classification (NCD). We can rename each of the 17 NCs using the shared features of integrative molecular and phenotypic profiles (SM section 7.4; Fig. S11c & Table S12, Data S11-14). For example, NC08 could be denoted as the “limbic system development-vision disorders-related diseases” since the most enriched PPI module (p=4.9e-324, Relevance ratio=0.7778) of its constituent diseases was mainly related to the GO biological process; “limbic system development” (p=1.13e-04), along with the fact that 73.84% (127/172) of diseases in NC08 shared the phenotype, “vision disorders” (p=4.9e-324) (Table S13-S14).

### New disease categories define diseases with clearer boundaries and balanced diversity

To confirm the quality of our overlapping disease categories, we compared the modularity of NCD with that of the ICD taxonomy. We found that the 17 NCs consistently have much higher modularity than the original ICD chapters for all types of disease association networks (SM section 7.3; Fig. 4a-h, Fig. S13). This finding indicates that the phenotypic and molecular links between the diseases of a category in NCD taxonomy are much closer than those in the ICD taxonomy.

**Figure 4.**
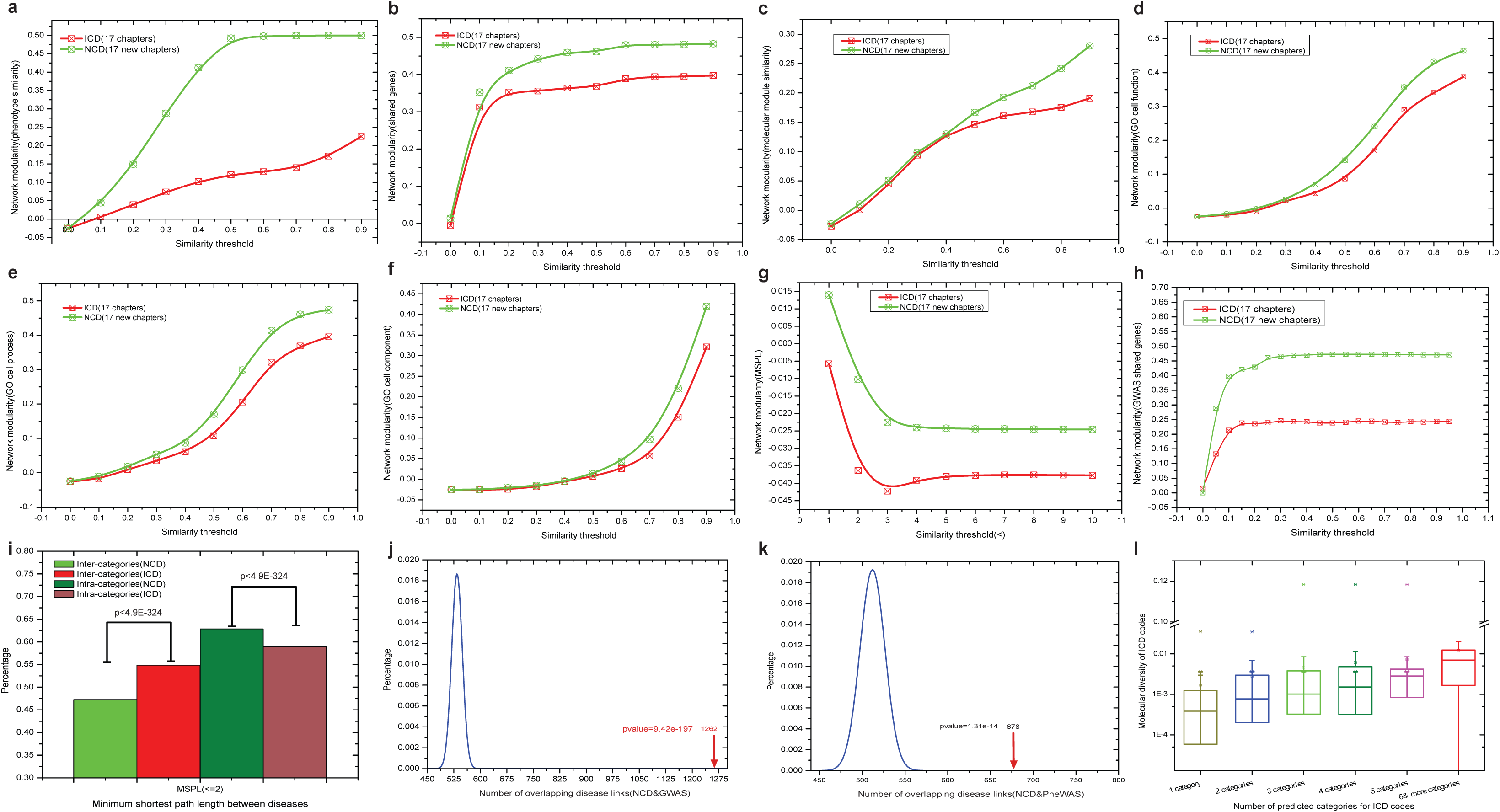
Properties of new disease categories (NCD) and comparison to conventional ICD classification. **a.** Modularity of phenotype(symptom)-based disease network (NCD vs. ICD); **b.** Modularity of gene-based disease network (NCD vs. ICD); **c.** Modularity of molecule module-based disease network (NCD vs. ICD); **d.** Modularity of gene ontology (molecular function)-based disease network (NCD vs. ICD); **e.** Modularity of gene ontology (biological process)-based disease network (NCD vs. ICD); **f.** Modularity of gene ontology (cellular component)-based disease network (NCD vs. ICD); **g.** Modularity of shortest path-based disease network (NCD vs. ICD); **h.** Modularity of GWAS shared gene-based disease network (NCD vs. ICD); **i.** Percentage of minimum shortest path lengths within the range [0, 2] of intra-category and inter-category (NCD vs. ICD, chi-squared test); **j.** The number of overlapping disease pairs from NCD with shared genes from GWAS, compared to random expectation, binomial test; **k.** The number of overlapping disease pairs from NCD with shared genes from PheWAS, compared to random expectation, binomial test; **l.** The number of predicted NCD categories for ICD codes (i.e. diseases) as a function of their molecular diversity.

Furthermore, we found that the minimum shortest path lengths (MSPLs) in PPI between disease pairs in the same NCD categories had a larger percentage of low values (i.e., [0,2]) than those in the ICD (Fig. 4i, 62.86% vs 58.95%, p<4.9e-324;SM section 7.3). This result indicates that diseases within an NCD category have a significantly higher degree of shared genes (or shorter path lengths) than diseases within a category in ICD. On the other hand, the MSPLs between disease pairs in different NCD categories had a significantly lower percentage of low values than those in the ICD (47.27% vs 54.88%, p<4.9e-324, Fig. 4i & Fig. S14; External validations in SM 8.3), which indicates a lower degree of shared genes (or shorter path lengths) between diseases from different categories in NCD than in ICD. These findings demonstrate that our NCD framework has clearer boundaries between distinct diseases belonging to different categories than those in the original ICD disease taxonomy. Moreover, to validate the robustness of NCD predictions, we calculated the degree of molecular associations among the diseases (in terms of network density) in each sub-category of NCD by investigating the overlaps with the disease pairs with shared genes from two independent phenotype-genotype association databases, namely GWAS and PheWAS (see Methods, SM section 1.2,1.6 & 8). We found that for the 223 sub-categories in NCD, network density was significantly higher compared to random controls (GWAS: p-value=9.42e-197, Fig. 4j;PheWAS:p-value=1.31e-14, Fig. 4k). This means that the diseases in the 223 sub-categories in NCD would tend to have a high degree of shared genes. For example, the New Chapter: NC12 in NCD, including 11 sub-categories and 136 ICD diseases (belonging to eight ICD chapters), is enriched with respiratory and airway diseases (e.g. COPD and asthma). We obtained 37 overlapped diseases from the GWAS database, which have a high degree of shared genes with the diseases in each sub-category of the NC12 (Fig. 5a). In particular, the sub-categories, such as NC12.M06 (p-value=2.53e-30), NC12.M03 (p-value=1.80e-38) and NC12.M02 (p-value=6.89e-19) have significantly higher density than those of the whole GWAS disease network (Fig. S19). Furthermore, we found that the overlapping subcategories of the NCD are able to differentiate between different components (i.e. asthma/allergy vs. COPD) of the same broad group of diseases (i.e. respiratory diseases) (see Fig. S19 for a detailed example). Indeed, in the NC12 disease chapter chiefly containing respiratory diseases, the two sub-categories, namely NC12.M06 and NC12.M07, overlap in the underlying molecular interaction network while still containing the respective disease (asthma and COPD, respectively) genes separately (Fig. 5a). A detailed discussion is offered in SM section 8(with results in Data S21-22, Table S18-19 & Fig. S20-S22).

**Figure 5.**
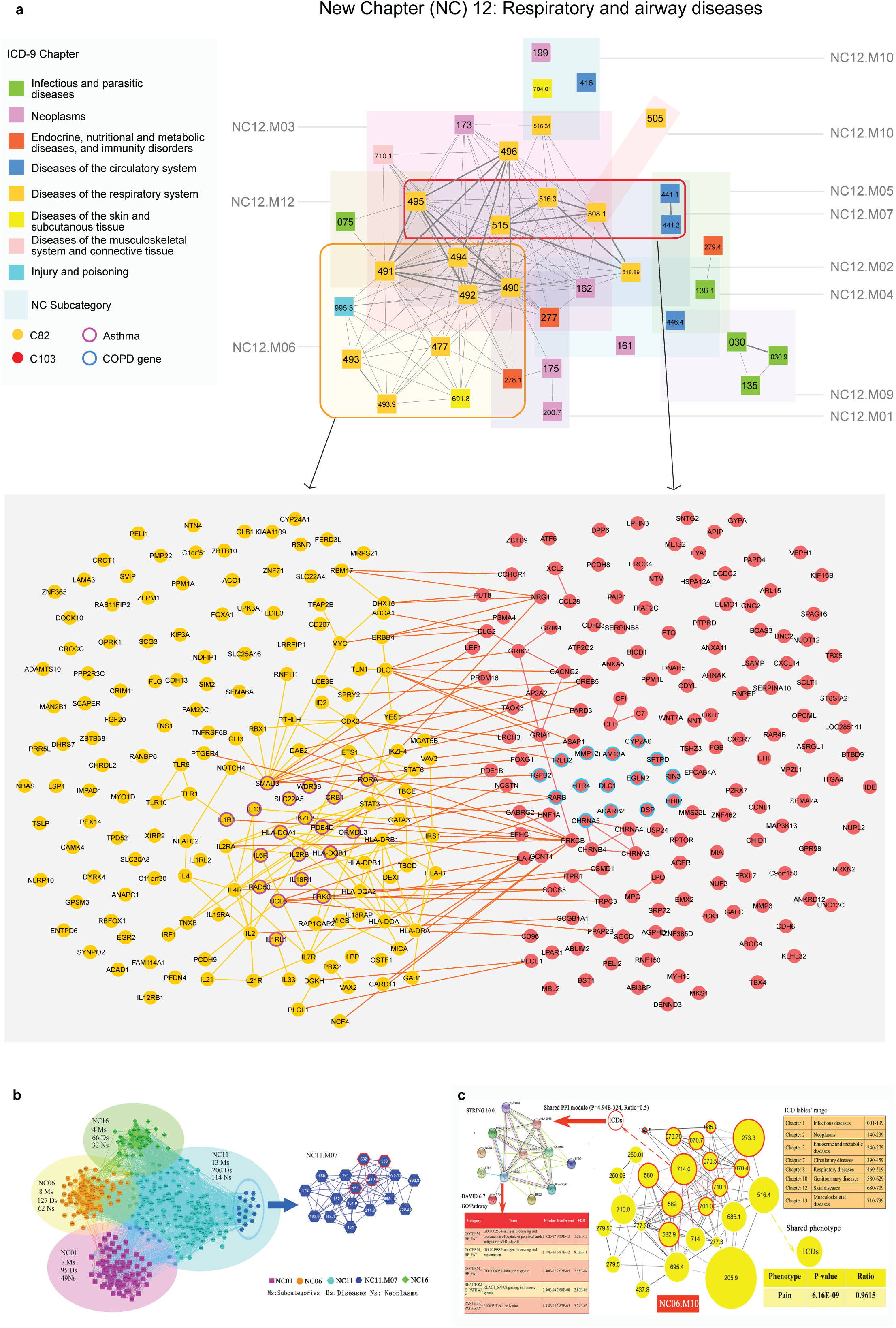
Biological insights of new disease taxonomy. **a.** The New Chapter containing airway diseases (NC12) consists of 11 sub-categories and 136 ICD diseases belonging to 8 ICD chapters. The subcategories overlap in the underlying molecular interaction network, while still separately including the disease genes (asthma and COPD, respectively) that characterize each subcategory; **b.** The disease network of neoplasms in NCD. The 32 sub-categories significantly representing neoplasms are divided into 4 NCs (G1). *Helicobacter pylori* [*H. pylori*] (041.86), malignant neoplasm of stomach (151), duodenal ulcer (532), peptic ulcer, site unspecified (533), which have significant relationships, are clearly clustered into a sub-category (NC11. M07) (G2); **c.** A sub-category (NC06.M10) in NCD, which includes diseases from 8 different ICD chapters with shared molecular mechanism and phenotypes. Fifty percent (13/26=50%) of the diseases in NC06.M10 share a PPI module, the biological function of which is enriched with immune system response, while over ninety percent (25/26=96.2%) of the shared common phenotype of this module is “Pain”.

In addition, in NCD, a disease can be classified into multiple categories, and the number of categories of a disease positively correlates with its molecular diversity (Fig. 4l, PCC=0.352, 95% CI=[0.311, 0.392], p-value < 4.94e-324; External validations in SM 8.3). For example, we reclassified neoplastic diseases into multiple categories due to their high molecular diversity. Two hundred and fifty-eight neoplastic diseases in our NCD were divided into 144 sub-categories and 17 NCs (Fig. S17 & S18). Thirty-nine out of 144 sub-categories (27.08%) were enriched with “neoplasm” diseases (Data S19, p-value = 2.78e-5). There were mainly 4 NCs (i.e., NC01, NC06, NC11, and NC16) containing these 32 sub-categories and 188 “neoplasm” disease codes (Fig. 5b), where 76.06% (143/188) of the neoplastic diseases were classified into more than 1 sub-category, ranging from 2 to 15(Data S20 & Table S17). The neoplasm with the highest MD, “malignant neoplasm of connective and other soft tissue” (ICD: 171; MD: 0.035), was reclassified into 15 sub-categories, and “malignant neoplasms of thyroid gland” (ICD: 193; MD: 0.0028) was assigned to 14 sub-categories. Furthermore, related diseases had been reclassified together in NCD, such as the well-known disease-correlations among *H. pylori* infection (ICD: 041.86), stomach cancer (ICD: 151), and duodenal ulcer (ICD: 532) or peptic ulcer (ICD: 533) (Fig. 5b) (Sitas 2016; Graham 2015).

More interestingly, some diseases, like viral hepatitis C (ICDs: 070.4, 070.5, 070.7), graft-versus-host disease (ICDs: 279.5/279.50), glomerulonephritis (ICDs: 580, 582, 582.9), circumscribed scleroderma (ICD: 701.0), systemic lupus erythematosus (ICD: 710.0), and rheumatoid arthritis (ICDs: 714/714.0), each from different chapters in ICD taxonomy, were classified together into a unique NCD sub-category (NC06.M10) since 50% (13/26) of these diseases share a PPI module related to immune response (SM section 7.4, Fig. 5c, Data S15-16, Table S15-16). In addition, diseases originally in the same ICD chapter, such as viral pneumonia (ICD: 480) and influenza (ICD: 487) from respiratory system-related diseases (Chapter 8), were reclassified into different categories in the NCD (NC12, NC10). Influenza shared more phenotype profiles with “episodic mood disorders” (ICD: 296) in NC10.M01, rather than viral pneumonia in NC12 (Fig. S15 & Data S17), which is in accordance with recent epidemiological studies between episodic mood disorders and influenza (Okusaga et al. 2011; Canetta et al. 2014), and, furthermore, we also found that influenza shared some molecular profiles with “episodic mood disorders” (ICD: 296) in NC10.M01 (Fig. S16, Data S18). These findings suggest that NCD offers a promising integrative framework incorporating both clinical phenotypes and molecular profiles for disease taxonomy that has very practical implications for the precise investigation of disease subtyping and etiologies.

## Discussion

Given the molecular network mechanisms (Barabasi, Gulbahce, and Loscalzo 2011; Zanzoni, Soler-Lopez, and Aloy 2009), genetic pleiotropy (Solovieff et al. 2013), as well as complicated genotype-phenotype associations underlying diseases, the establishment of a molecular-based disease taxonomy with clear boundaries is essential but challenging. From the molecular network perspective, we first investigated the utility, shortcomings, and inconsistencies of ICD-9-CM, the established disease taxonomy for clinical settings. We found that there exist a considerable number (~40% of our investigated diseases) of diseases, for example, cancer and infectious diseases that have diverse molecular network mechanisms and tend to interact more with diseases from other chapters. It is also these molecularly diverse diseases that mainly contribute to the blurred boundary of ICD disease taxonomy (see Methods, SM section 4 & 7). As a result of exploring the molecular diversity and cross-chapter interactions between diseases, we propose a novel disease classification system based on the integration of the clinical phenomic and molecular profiles of diseases. In particular, we integrate disease networks taking into account molecular and phenotypic connectivity among diseases, predict the multiple disease categories that diseases belong to, and finally validate the biological cohesiveness of our NCD by network topological measures such as modularity and shortest path length. Our findings indicate that although general correlations exist between disease closeness in ICD taxonomy and underlying molecular profiles, ICD still displays significant limitations with regard to the heterogeneity of molecular diversity and clear category boundaries. In our NCD, a disease with a high molecular diversity tends to be classified into multiple disease categories, which indicates that there exist more disease subtypes for that disease. For example, “malignant neoplasm of the pancreas” was reclassified into 11 sub-categories and 4 NCs, which is consistent with a recent study wherein 4 phenotypic subtypes of pancreatic cancer were enriched for 10 distinct molecular mechanisms (Bailey et al. 2016). Therefore, we believe that the new disease classification system may help facilitate precise clinical diagnosis and correct prognosis (Jameson and Longo 2015), and does so in alignment with refined molecular network diagnostics. Furthermore, the molecular network underpinnings and overlapping disease categories of NCD provide a credible relationship map between diseases and disease categories that may radically transform our current understanding of diseases and relevant treatment paradigms. On the one hand, our approach accurately links diseases with all possible underlying mechanisms in the molecular interaction network. On the other hand, it presents a promising approach to the identification of targeted drugs for the treatment of related diseases. For example, breast cancer and influenza (both in NC11.M02) may share potential drug targets (Park 2012). As another example, metformin, widely prescribed to treat metabolic syndrome (in NC11.M02), could alter the gut microbiome composition and function, improve gut microbial dysbiosis (Forslund et al. 2015; Cabreiro et al. 2013), and also prevent colorectal cancer (also in NC11.M02) through microbiome-influenced immune response modification (Nakatsu et al. 2015). Here, it is important to note that while a considerable number of diseases have a strong environmental component, here our main focus has been the many diverse molecular determinants. In the future, additional environmental factors such as epigenetic changes can be added into the data integration scheme to further refine the classification.

A limitation of this analysis is that DiseaseConnect yields an incomplete disease-gene database (Menche et al. 2015): only 1,883 ICD diseases could be mapped (Table S11), leading to only 1,797 diseases included in the NCD. Additionally, our NCD merely delivers a two-level taxonomy framework without elaborated hierarchical structures in the same disease categories, which could be further refined or optimized through methods like hierarchical clustering algorithms (Murtagh and Contreras 2012). In this big-data era, the dramatically increasing multi-omics databases, as well as clinical data from electronic health records (EHR) involving phenotypic, therapeutic and environmental factors information (Jensen, Jensen, and Brunak 2012), should also be incorporated into the new disease taxonomy refinement for patient stratification and disease treatment. At this point, a realistic assumption is that the translation of this classification to the clinic will need some time. That said, while the ICD is originally made “by clinicians for clinicians”, it is now widely used by biomedical researchers as well to gain a deeper understanding of human diseases. We therefore believe that researchers will be the first and direct beneficiaries of our approach.

In conclusion, our study provides valuable insights into the polyhierarchical network-based disease classification beyond the traditional tree structure. Our integrated disease network approach is sufficiently powerful to elucidate the tangled underpinnings of human diseases and uncover distinct disease boundaries. Our work may provide a new framework for the disease taxonomy reform based on big-data fusion, so as to generate further the robust infrastructure needed for precision medicine.

## Methods

### Basic datasets compilation

In this work, large curation efforts are performed to generate the related data sources (details see SM section 1). We obtained the updated text version of ICD-9-CM (2011) and extracted the list of ICD codes with their hierarchical structures. While we recognize the improvements of the currently used ICD-10 over ICD-9, nevertheless, we chose to use ICD-9-CM as the adoption of ICD-10 has been slow in the United States(Butler 2014) and since it was still being widely used at the time of the data collection for this paper(Blair et al. 2013; Wang et al. 2017). Furthermore, although ICD-10 does have more codes than ICD-9-CM, the structure is kept almost the same. We obtained the high-quality phenotype-genotype associations from DiseaseConnect database (2015 version), leaving out the less reliable text mining entries and focusing only on GWAS, OMIM and differential expression evidence types, and manually mapped those diseases to ICD codes. To calculate the molecular network and phenotype characteristics related to disease phenotypes, a high-quality subset of human protein-protein interactions was filtered from STRING V9.0 using the score threshold at >=700, as well as a well-established disease-phenotype association dataset (i.e. HSDN) derived from PubMed bibliographic records and the gene ontology annotations from NCBI gene database are adopted. While we chose to use STRING, which is a widely used protein-protein interaction database, to ensure the results are not biased by computational predictions that are not as reliable as experimental ones, we have repeated the classification pipeline with manually curated PPI networks (Menche et al. 2015) with only experimental results and found that the results are robust (SM section 8.3). In addition, to validate the robustness of our results from independent data sources, we filtered the GWAS and PheWAS data from UCSC Genome Browser(Tyner et al. 2017) and PheWAS catalog(Denny et al. 2010) respectively, and performed additional ICD mapping task to prepare the data for validation analysis. The GWAS evidence of the DiseaseConnect database, which we used to build the disease associations, comes from the NHGRI GWAS catalog(Welter et al. 2014), whereas for validation, we used the UCSC-GWAS Genome Browser. We have ensured that the GWAS data used to build the networks and to validate them have a very small overlap (SM Section 8).

### Evaluating the quality of ICD disease taxonomy

Here, we systematically evaluated the consistency of disease categories in ICD taxonomy from both clinical phenotype and molecular profiles (details are in SM section 2). We investigated the quality of ICD disease taxonomy by evaluating the correlation between the closeness of disease pairs in the disease taxonomy and their underlying molecular connections (and phenotype similarities). For example, if two disease pairs have close positions (e.g. have a low level common parent disease) in the disease taxonomy, then we would expect that those disease pairs might have common genes or shared protein-protein interactions or similar phenotypes. We calculated the category similarity between disease pairs using a widely used semantic similarity measure (i.e. Lin measure using information content) to represent the closeness of disease pairs located in the ICD taxonomy. The molecular and phenotype similarity between disease pairs are calculated from the perspectives of shared genes and their GO annotations, network pathologies and shared phenotypes according to well established similarity measures (e.g. Cosine measure and Jaccard measure). In particular, to propose a more robust representation of genetic profiles of diseases, we partitioned the STRING network into 314 topological modules (Data S2) and used them to construct the relevant module vectors of diseases using Odds Ratio (OR) as weighting measure. For example, an ICD disease code would be represented with a 314-dimensional vector, which has a value of w_ij_ if its related gene is in a module or 0 otherwise. Suppose we have N genes in total and m_i_ genes of a module i. Now for a disease d_i_ with n_j_ genes, which has k_ij_ overlapping genes with the module i, we calculated the value of w_ij_ as the following equation,

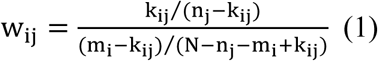

We used the cosine measure to calculate the molecular module similarity between disease pairs after the molecular module vector (i.e. OR weighting) of each disease was constructed.

Furthermore, if ICD taxonomy proposes a good category framework for organizing the diseases, there would exist much more molecular interactions or phenotype relationships between the diseases of the same chapters than those of the different chapters. Therefore, when we consider the ICD chapters as the module annotations (i.e. all the diseases in one chapter would be considered as members of a same module) for disease association networks (the disease networks with molecular or phenotype associations as links), the modularity of the disease association network could reflect the quality of ICD disease taxonomy. This means that the higher the modularity, the higher the quality of the ICD chapters as a disease category framework.

To evaluate the quality of community structures in complex network, the modularity measure (Newman 2006) was proposed to quantify the extent to which the connection in communities is above the random expectation in the whole network. Let a network have m edges and A_vw_ be an element of the adjacency matrix of the network. Suppose the vertices in the network are divided into communities such that vertex v belongs to community c_v_. Then the modularity Q is defined as:

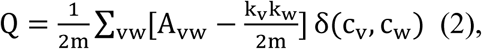

where the function δ(i, j)is 1 if i = j and 0 otherwise, and k_v_ is the degree of vertex v. The value of the modularity lies in the range [−1/2,1]. It is positive if the number of edges within groups exceeds the number expected on the basis of chance. Otherwise, it would be negative. We use it to measure the consistency of disease categories (ICD chapter or NCD) as an annotation of topological module (or community) structures within disease networks. We suppose that if a disease category framework is good enough from the molecule or phenotype profile perspective, then there would be more links existing between the disease members in a category than random expectation.

### Measuring the disease specificity

As a quantification of the molecular diversity (or the inverse specificity) of a disease, we calculated the maximum betweenness of disease-related genes in the PPI network (Data S3). Betweenness^(Freeman 1977)^ is a widely used centrality measure to quantify how many shortest paths run through a given node. In particular, bridging nodes that connect disparate components of the network often have a high betweenness. The betweenness centrality of a node v is given by:

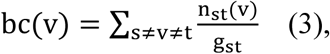

where n_st_(v)denotes the number of shortest paths from sto t that pass through v and g_st_ is the total number of shortest paths from s to t. We will adopt the convention that 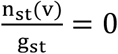 if both n_st_(v) and g_st_ are zero. This means that we assume the molecular diversity of diseases would largely lie on the related genes with maximum betweenness. For example, to quantify the molecular diversity (in terms of maximum betweenness) of Alzheimer’s disease (AD), we calculated all the betweenness values for the AD-related genes, such as APP, APOE, TNF and NOS3. Finally, we considered the molecular diversity of AD as 8.44e-3 since we found that APP has the maximum betweenness of 8.44e-3 among those genes (see Fig. S5a). In fact, this kind of measurement has been successfully used in a previous study(Zhou et al. 2014) to evaluate the diversity of diseases, which indicated that the diversity of disease manifestations has a strong positive correlation with the molecular diversity of diseases. For disease taxonomy with good quality, we would expect it to have its lowest level diseases (the leaf nodes in the tree-structure disease taxonomy) with similar molecular diversities.

### Detection of the significant disease-chapter associations

We calculated the edge density to quantify the molecular interactions between ICD chapters. To further detect the significant interactions between diseases in different chapters, we find an approach to obtain the diseases that have significant interactions with diseases in chapters other than their own. Given a disease d_i_ for investigation, we evaluate whether the proportion of interactions (i.e. edge density) of d_i_ to the disease set D_ck_ of a chapter C_k_ is significantly larger than the average proportion of interactions between the diseases in C_k_ (Fig. S6). We use binomial test to filter the significant interacting disease-chapter pairs, in which the edge density of the disease to the chapter is significantly higher than the average edge density of the diseases in the corresponding chapter (details are in SM section 4).

### Multi-category prediction of diseases

The results showing positive correlations between CS and molecular similarity, and the high MD of many diseases imply that it would be possible to predict the multi-category map for each disease using its underlying molecular connections. To demonstrate a pilot method for multiple disease category prediction by integrating molecular module and shared gene similarities, we provided a novel algorithm to generate the possible associated additional disease categories for a given disease with the corresponding molecular association scores. (details are in SM section 5, Fig. S7). In this algorithm, we integrated the correlation between category similarity and module similarity with significant disease-chapter associations (which are based on the shared gene similarity) to predict the additional chapters for a given disease. We divide the disease pairs in the same chapter to three subsets, which correspond to those pairs with shared root parents, shared second-level intermediate parents and shared third-level intermediate parents, respectively, to help predict to what degree a pair of diseases would be located closely in the disease taxonomy. The principle of the algorithm adheres to the positive correlation between category similarity (or the closeness of position of the disease pairs in ICD disease taxonomy) and molecular profile similarity of disease pairs, which means that strong molecular profile similarity between disease pairs would indicate close locations of them in the disease taxonomy. To ensure detecting the significant disease-chapter associations, we next filtered the predicted disease-chapter associations with positive association scores by the significant disease-chapter interactions based on shared genes.

### Construction of integrated disease network

To integrate disease associations derived from both molecular and phenotype features, we performed several sequential analytical steps to generate a high reliable disease network with strict filtering criterions of the disease links (details are in SM section 6). Firstly, we generated three disease association networks (i.e. SGDN with 133,469 links and 1868 nodes, HSDN with 1,639,791 links and 1814 nodes and MSDN with 598,420 links and 1744 nodes, Fig. S10 & Table S10) according to shared genes, shared phenotypes and molecular module similarity, respectively. To reduce the possible noise and bias of disease related data sources, we applied a multi-scale backbone algorithm(Serrano, Boguna, and Vespignani 2009) to obtain high reliable disease links (with significantly high weights than the random expectations) from the three disease networks. We finally obtained 53,241, 8,554 and 134,370 high reliable links for MSDN, SGDN and HSDN, respectively and retained most nodes (1,744 for MSDN, 1,782 for SGDN and 1,814 for HSDN) of these networks. To further reduce the possible weak associations (the disease pairs with high module similarity but no direct protein interactions) derived from module similarity, we calculated the MSPLs between each disease pairs and used it as a filtering criterion (with MSPL<=1) for MSDN, which resulted in a more biological meaningful subset of MSDN with 33,611 links and 1,694 diseases.

SGDN would capture high associations between disease pairs if they have high degree of shared genes even their related genes are not forming genetic modules. However, MSDN would give high weights for disease links if the disease pairs have similar co-locations on the topological modules of molecular network even they have no shared genes. Therefore, MSDN and SGDN are actually two complementary molecular association evidences for disease pairs and we finally obtained the union of the subset of MSDN and SGDN as the molecular association disease network (MADN), which contains 35,389 links and 1,811 nodes with the weights derived from the two original networks. Next, we adopted a highly strict criterion to obtain an integrated disease network (IDN) from the fusion of MADN and HSDN links, which contains 35,114 disease links and 1,857 nodes.

### Overlapping category detection from integrated disease network

Finding the overlapping disease categories could be transformed to the task of detecting the overlapping communities (i.e. modules) from the IDN. BigClam(Yang and Leskovec 2013) is a state-of-the-art overlapping community detection algorithm based on a variant of non-negative matrix factorization, which achieves near linear running time and comparable high quality community results. We used the BigClam algorithm, which is packaged in SNAP complex network software (http://snap.stanford.edu/snap/) to automatically detect overlapping communities from IDN network. Finally, we obtained 223 overlapping disease communities with 1,797 distinct ICD disease codes. These 223 disease subcategories contain different numbers of ICD codes, ranging from 5 to 168 (Fig. S12 & Data S10).

To obtain a top-level category framework of diseases corresponding to the chapters in ICD taxonomy, we calculated the overlapping degree of the 223 disease sub-categories by using Jaccard similarity to measure the common number of diseases held by two given disease categories. This generated a disease category network with 2,685 links representing shared ICD codes (a link is established if two disease categories share at least an ICD code and the weights of links correspond to the Jaccard similarity) and nodes representing disease categories. After that, we clustered the 223 disease sub-categories additionally by a widely used non-overlapping community detection algorithm (considering the link weight and setting the resolution parameter as 0.5) into 17 top-level categories (which corresponds to the number of original chapter-level categories in ICD, which we named as New Chapters, NCs) using the shared ICD codes (Fig. S11c & Data S10). The modularity of these 17 top-level categories (this makes a good comparable partition with ICD chapters) in the network of 223 sub-categories is 0.426, which means a rather good partition of the network. These 17 NCs contain different numbers of sub-categories ranging from 4 to 25 or of diseases ranging from 53 to 369 (Fig. S11c & Data S10), covering diseases from all of the 17 chapters of ICD taxonomy (Table S11). These 17 NCs would still contain overlapping disease codes since the 223 disease sub-categories have overlapping disease codes. Therefore, 17 NCs with 223 disease sub-categories form a disease taxonomy consisting of two hierarchical levels with polyhierarchical categories although with a limited number (1,797) of disease members.

### Statistical validation of NCD from external data

To validate the robustness of NCD, we obtained two external phenotype-genotype data sources (i.e. UCSC-GWAS and PheWAS catalog), which have not been integrated yet for generating NCD for further investigation. By measuring whether the disease members in the sub-categories in NCD tend to incorporate the associations of shared genes from these two data sources, we would be able to validate the quality of NCD. If the diseases in such NCD sub-categories would tend to involve shared genes, then the diseases would be more likely associated with one another than other diseases. To test this hypothesis, we obtained the overlapping disease codes (ODC) in both NCD and the two external phenotype-genotype association databases and evaluate the degree of these ODC disease links in each NCD sub-category when considering two diseases linked if they share common genes. In detail, we firstly obtained the common disease codes involved in both NCD and UCSC-GWAS or PheWAS database. Then we generated a disease network with shared genes derived from the two datasets, in which two diseases linked if they shared at least one common gene. After that for each NCD sub-category, we generated a complete disease network with the ODC diseases in it and overlaid the network on the disease network with shared genes. Finally, the overlapping percentage of disease links would be calculated for evaluating the degree of molecular associations involved in diseases in each NCD sub-categories (details are in SM section 8, Fig. S20-S22).

### Statistical analysis

We use R 3.1.0 as the main statistical tool in our work. The comparison of two percentages was calculated by Binomial test or Chi-squared test. Wilcoxon rank sum test was used for compare two independent list of values (e.g. two types of molecular diversities and two groups of MSPLs). All the correlations between two variables were calculated by Pearson’s product moment correlation coefficient. Due to the incompleteness and bias of disease-related data (i.e. disease-gene associations and disease-symptom associations), we need to distinguish the information from the background noise. Therefore, for comparison with random expectation, we reshuffle (100 random permutations) the symptom features and the related genes of each disease using the Fisher-Yates method (Fisher and Yates 1948). The calculations from random permutations were used for the correlation between CS and molecular similarity, as well as phenotype similarity. In addition, this was used for detection of the disease categories with high molecule diversity.

## Author Contributions

Z.W., A.S., X.Z. and J.L.(Joseph Loscalzo) conceived and designed the research; X.Z., L.L., J.L.(Jun Liu), A.H., Z.Y., B.L., Z.G., L.G., C.S., J.L.(Joseph Loscalzo) and Z.W. performed the research tasks: data curation and compiling(L.L., J.L.(Jun Liu), Z.Y., B.L. and Z.G.),data analysis(X.Z. and G.L.), result validation(L.L.,J.L.(Jun Liu),C.S., A.H., J.L.(Joseph Loscalzo) and Z.W.); X.Z, J.L.(Jun Liu), A.H., A.S. and L.L. wrote the manuscript. All authors have reviewed and revised the manuscript.

## Acknowledgments

The work was supported by National Natural Science Foundation of China (61105055, 81230086 and 81673833), China 863 Program (2003AA2Z2022) and the Fundamental Research Funds for the Central public welfare research institutes (ZZ0908029). We also acknowledge the support by National Institutes of Health (NIH) grants P50-533 HG004233-CEGS, MapGen grant (U01HL108630) and P01 HL083069, U01 534 HL065899, P01 HL105339, R01HL111759, 1P01HL132825-01 and RC HL10154301.

## References

Ahmad, T. S. Marshall, and D. Jewell. 2003. ‘Genotype-­‐based phenotyping heralds a new taxonomy for inflammatory bowel disease’, Curr Opin Gastroenterol 19: 327–35.

Alizadeh, A. A., M. B. Eisen, R. E. Davis, C. Ma, I. S. Lossos, A. Rosenwald, J. C. Boldrick, H. Sabet, T. Tran, X. Yu, J. I. Powell, L. Yang, G. E. Marti, T. Moore, J. Hudson, Jr., L. Lu, D. B. Lewis, R. Tibshirani, G. Sherlock, W. C. Chan, T. C. Greiner, D. D. Weisenburger, J. O. Armitage, R. Warnke, R. Levy, W. Wilson, M. R. Grever, J. C. Byrd, D. Botstein, P. O. Brown, and L. M. Staudt. 2000. ‘Distinct types of diffuse large B-cell lymphoma identified by gene expression profiling’, Nature, 403: 503–11.

Arostegui, I., C. Esteban, S. Garcia-Gutierrez, M. Bare, N. Fernandez-de-Larrea, E. Briones, and J. M. Quintana. 2014. ‘Subtypes of patients experiencing exacerbations of COPD and associations with outcomes’, PLoS One, 9: e98580.

Ashburn, T. T., and K. B. Thor. 2004. ‘Drug repositioning: identifying and developing new uses for existing drugs’, Nat Rev Drug Discov, 3: 673–83.

Bailey, P., D. K. Chang, K. Nones, A. L. Johns, A. M. Patch, M. C. Gingras, D. K. Miller, A. N. Christ, T. J. Bruxner, M. C. Quinn, C. Nourse, L. C. Murtaugh, I. Harliwong, S. Idrisoglu, S. Manning, E. Nourbakhsh, S. Wani, L. Fink, O. Holmes, V. Chin, M. J. Anderson, S. Kazakoff, C. Leonard, F. Newell, N. Waddell, S. Wood, Q. Xu, P. J. Wilson, N. Cloonan, K. S. Kassahn, D. Taylor, K. Quek, A. Robertson, L. Pantano, L. Mincarelli, L. N. Sanchez, L. Evers, J. Wu, M. Pinese, M. J. Cowley, M. D. Jones, E. K. Colvin, A. M. Nagrial, E. S. Humphrey, L. A. Chantrill, A. Mawson, J. Humphris, A. Chou, M. Pajic, C. J. Scarlett, A. V. Pinho, M. Giry-Laterriere, I. Rooman, J. S. Samra, J. G. Kench, J. A. Lovell, N. D. Merrett, C. W. Toon, K. Epari, N. Q. Nguyen, A. Barbour, N. Zeps, K. Moran-Jones, N. B. Jamieson, J. S. Graham, F. Duthie, K. Oien, J. Hair, R. Grutzmann, A. Maitra, C. A. Iacobuzio-Donahue, C. L. Wolfgang, R. A. Morgan, R. T. Lawlor, V. Corbo, C. Bassi, B. Rusev, P. Capelli, R. Salvia, G. Tortora, D. Mukhopadhyay, G. M. Petersen, D. M. Munzy, W. E. Fisher, S. A. Karim, J. R. Eshleman, R. H. Hruban, C. Pilarsky, J. P. Morton, O. J. Sansom, A. Scarpa, E. A. Musgrove, U. M. Bailey, O. Hofmann, R. L. Sutherland, D. A. Wheeler, A. J. Gill, R. A. Gibbs, J. V. Pearson, A. V. Biankin, and S. M. Grimmond. 2016. ‘Genomic analyses identify molecular subtypes of pancreatic cancer’, Nature, 531: 47–52.

Barabasi, A. L., N. Gulbahce, and J. Loscalzo. 2011. ‘Network medicine: a network-based approach to human disease’, Nat Rev Genet, 12: 56–68.

Bianchini, G., J. M. Balko, I. A. Mayer, M. E. Sanders, and L. Gianni. 2016. ‘Triple-negative breast cancer: challenges and opportunities of a heterogeneous disease’, Nat Rev Clin Oncol, 13: 674–90.

Blair, D. R., C. S. Lyttle, J. M. Mortensen, C. F. Bearden, A. B. Jensen, H. Khiabanian, R. Melamed, R. Rabadan, E. V. Bernstam, S. Brunak, L. J. Jensen, D. Nicolae, N. H. Shah, R. L. Grossman, N. J. Cox, K. P. White, and A. Rzhetsky. 2013. ‘A nondegenerate code of deleterious variants in Mendelian loci contributes to complex disease risk’, Cell, 155: 70–80.

Butler, M. 2014. ‘Not so fast! Congress delays ICD-10-CM/PCS. Examining how the delay happen, its industry impact, and how best to proceed’, J AHIMA, 85: 24–8.

Cabreiro, F., C. Au, K. Y. Leung, N. Vergara-Irigaray, H. M. Cocheme, T. Noori, D. Weinkove, E. Schuster, N. D. Greene, and D. Gems. 2013. ‘Metformin retards aging in C. elegans by altering microbial folate and methionine metabolism’, Cell, 153: 228–39.

Cancer Genome Atlas Research, Network, University Analysis Working Group: Asan, B. C. Cancer Agency, Brigham, Hospital Women’s, Institute Broad, University Brown, University Case Western Reserve, Institute Dana-Farber Cancer, University Duke, Centre Greater Poland Cancer, School Harvard Medical, Biology Institute for Systems, K. U. Leuven, Clinic Mayo, Center Memorial Sloan Kettering Cancer, Institute National Cancer, Hospital Nationwide Children’s, University Stanford, Alabama University of, Michigan University of, Carolina University of North, Pittsburgh University of, Rochester University of, California University of Southern, M. D. Anderson Cancer Center University of Texas, Washington University of, Institute Van Andel Research, University Vanderbilt, University Washington, Institute Genome Sequencing Center: Broad, Louis Washington University in St, B. C. Cancer Agency Genome Characterization Centers, Institute Broad, School Harvard Medical, University Sidney Kimmel Comprehensive Cancer Center at Johns Hopkins, Carolina University of North, Center University of Southern California Epigenome, M. D. Anderson Cancer Center University of Texas, Institute Van Andel Research, Institute Genome Data Analysis Centers: Broad, University Brown, School Harvard Medical, Biology Institute for Systems, Center Memorial Sloan Kettering Cancer, Cruz University of California Santa, M. D. Anderson Cancer Center University of Texas, Consortium Biospecimen Core Resource: International Genomics, Hospital Research Institute at Nationwide Children’s, Services Tissue Source Sites: Analytic Biologic, Center Asan Medical, Bioscience Asterand, Hospital Barretos Cancer, BioreclamationIvt, Clinic Botkin Municipal, School Chonnam National University Medical, System Christiana Care Health, Cureline, University Duke, University Emory, University Erasmus, Medicine Indiana University School of, Moldova Institute of Oncology of, Consortium International Genomics, Invidumed, Hamburg Israelitisches Krankenhaus, Medicine Keimyung University School of, Center Memorial Sloan Kettering Cancer, Goyang National Cancer Center, Bank Ontario Tumour, Centre Peter MacCallum Cancer, School Pusan National University Medical, School Ribeirao Preto Medical, Hospital St. Joseph’s, Center Medical, University St. Petersburg Academic, Bank Tayside Tissue, Dundee University of, Center University of Kansas Medical, Michigan University of, Hill University of North Carolina at Chapel, Medicine University of Pittsburgh School of, M. D. Anderson Cancer Center University of Texas, University Disease Working Group: Duke, Center Memorial Sloan Kettering Cancer, Institute National Cancer, M. D. Anderson Cancer Center University of Texas, Medicine Yonsei University College of, Csra Inc Data Coordination Center, and Health Project Team: National Institutes of. 2017. ‘Integrated genomic characterization of oesophageal carcinoma’, Nature, 541: 169–75.

Canetta, S. E., Y. Bao, M. D. Co, F. A. Ennis, J. Cruz, M. Terajima, L. Shen, C. Kellendonk, C. A. Schaefer, and A. S. Brown. 2014. ‘Serological documentation of maternal influenza exposure and bipolar disorder in adult offspring’, Am J Psychiatry, 171: 557–63.

Chong, C. R., and D. J. Sullivan, Jr. 2007. ‘New uses for old drugs’, Nature, 448: 645–6.

Chuang, H. Y., E. Lee, Y. T. Liu, D. Lee, and T. Ideker. 2007. ‘Network-based classification of breast cancer metastasis’, Mol Syst Biol, 3: 140.

Cimino, J. J. 2011. ‘High-quality, standard, controlled healthcare terminologies come of age’, Methods Inf Med, 50: 101–4.

Council, National Research, Committee, Framework, Developing, New Taxonomy, and Disease (ed.)^(eds.). 2011. Toward Precision Medicine: Building a Knowledge Network for Biomedical Research and a New Taxonomy of Disease (The National Academies Press: Washington, DC).

Dahlem, D., D. Maniloff, and C. Ratti. 2015. ‘Predictability Bounds of Electronic Health Records’, Sci Rep, 5: 11865.

Denny, J. C., M. D. Ritchie, M. A. Basford, J. M. Pulley, L. Bastarache, K. Brown-Gentry, D. Wang, D. R. Masys, D. M. Roden, and D. C. Crawford. 2010. ‘PheWAS: demonstrating the feasibility of a phenome-wide scan to discover gene-disease associations’, Bioinformatics, 26: 1205–10.

Dienstmann, R., L. Vermeulen, J. Guinney, S. Kopetz, S. Tejpar, and J. Tabernero. 2017. ‘Consensus molecular subtypes and the evolution of precision medicine in colorectal cancer’, Nat Rev Cancer, 17: 79–92.

Evans, J. M., L. A. Donnelly, A. M. Emslie-Smith, D. R. Alessi, and A. D. Morris. 2005. ‘Metformin and reduced risk of cancer in diabetic patients’, BMJ, 330: 1304–5.

Fisher, Ronald A., and Frank Yates. 1948. Statistical tables for biological, agricultural and medical research (London: Oliver & Boyd.).

Forslund, K., F. Hildebrand, T. Nielsen, G. Falony, E. Le Chatelier, S. Sunagawa, E. Prifti, S. Vieira-Silva, V. Gudmundsdottir, H. Krogh Pedersen, M. Arumugam, K. Kristiansen, A. Y. Voigt, H. Vestergaard, R. Hercog, P. Igor Costea, J. R. Kultima, J. Li, T. Jorgensen, F. Levenez, J. Dore, H. B. Nielsen, S. Brunak, J. Raes, T. Hansen, J. Wang, S. D. Ehrlich, P. Bork, and O. Pedersen. 2015. ‘Disentangling type 2 diabetes and metformin treatment signatures in the human gut microbiota’, Nature, 528: 262–6.

Franceschini, A., D. Szklarczyk, S. Frankild, M. Kuhn, M. Simonovic, A. Roth, J. Lin, P. Minguez, P. Bork, C. von Mering, and L. J. Jensen. 2013. ‘STRING v9.1: protein-protein interaction networks, with increased coverage and integration’, Nucleic Acids Res, 41: D808–15.

Freeman, L C. 1977. ‘A set of measures of centrality based on betweenness’, Sociometry: 35–41.

Gligorijevic, V., N. Malod-Dognin, and N. Przulj. 2016. ‘Integrative methods for analyzing big data in precision medicine’, Proteomics, 16: 741–58.

Gligorijevic, V., and N. Przulj. 2015. ‘Methods for biological data integration: perspectives and challenges’, J R Soc Interface, 12.

Goh, K. I., M. E. Cusick, D. Valle, B. Childs, M. Vidal, and A. L. Barabasi. 2007. ‘The human disease network’, Proc Natl Acad Sci U S A, 104: 8685–90.

Golub, T. R., D. K. Slonim, P. Tamayo, C. Huard, M. Gaasenbeek, J. P. Mesirov, H. Coller, M. L. Loh, J. R. Downing, M. A. Caligiuri, C. D. Bloomfield, and E. S. Lander. 1999. ‘Molecular classification of cancer: class discovery and class prediction by gene expression monitoring’, Science, 286: 531–7.

Graham, D. Y. 2015. ‘Helicobacter pylori update: gastric cancer, reliable therapy, and possible benefits’, Gastroenterology Cell, 158: 719–31e3.

Grainge, C., P. S. Thomas, J. C. Mak, M. J. Benton, T. K. Lim, and F. W. Ko. 2016. ‘Year in review 2015: Asthma and chronic obstructive pulmonary disease’, Respirology, 21: 765–75.

Hidalgo, C. A., N. Blumm, A. L. Barabasi, and N. A. Christakis. 2009. ‘A dynamic network approach for the study of human phenotypes’, PLoS Comput Biol, 5: e1000353.

Hoadley, K. A., C. Yau, D. M. Wolf, A. D. Cherniack, D. Tamborero, S. Ng, M. D. Leiserson, B. Niu, M.D. McLellan, V. Uzunangelov, J. Zhang, C. Kandoth, R. Akbani, H. Shen, L. Omberg, A. Chu, A. A. Margolin, L. J. Van’t Veer, N. Lopez-Bigas, P. W. Laird, B. J. Raphael, L. Ding, A. G. Robertson, L. A. Byers, G. B. Mills, J. N. Weinstein, C. Van Waes, Z. Chen, E. A. Collisson, C. C. Benz, C. M. Perou, and J. M. Stuart. 2014. ‘Multiplatform analysis of 12 cancer types reveals molecular classification within and across tissues of origin’, Cell, 158, 929–44.

Hofmann-Apitius, M., M. E. Alarcon-Riquelme, C. Chamberlain, and D. McHale. 2015. ‘Towards the taxonomy of human disease’, Nat Rev Drug Discov, 14: 75–6.

Hofree, M., J. P. Shen, H. Carter, A. Gross, and T. Ideker. 2013. ‘Network-based stratification of tumor mutations’, Nat Methods, 10: 1108–15.

Hu, J. X., C. E. Thomas, and S. Brunak. 2016. ‘Network biology concepts in complex disease comorbidities’, Nat Rev Genet, 17: 615–29.

Jameson, J. L., and D. L. Longo. 2015. ‘Precision medicine-personalized, problematic, and promising’, N Engl J Med, 372: 2229–34.

Jensen, A. B., P. L. Moseley, T. I. Oprea, S. G. Ellesoe, R. Eriksson, H. Schmock, P. B. Jensen, L. J. Jensen, and S. Brunak. 2014. ‘Temporal disease trajectories condensed from population-wide registry data covering 6.2 million patients’, Nat Commun, 5: 4022.

Jensen, P. B., L. J. Jensen, and S. Brunak. 2012. ‘Mining electronic health records: towards better research applications and clinical care’, Nat Rev Genet, 13: 395–405.

Jeste, S. S., and D. H. Geschwind. 2014. ‘Disentangling the heterogeneity of autism spectrum disorder through genetic findings’, Nat Rev Neurol, 10: 74–81.

Lee, D. S., J. Park, K. A. Kay, N. A. Christakis, Z. N. Oltvai, and A. L. Barabasi. 2008. ‘The implications of human metabolic network topology for disease comorbidity’, Proc Natl Acad Sci U S A, 105: 9880–5.

Li, Y. Y., and S. J. Jones. 2012. ‘Drug repositioning for personalized medicine’, Genome Med, 4: 27.

Liu, C. C., Y. T. Tseng, W. Li, C. Y. Wu, I. Mayzus, A. Rzhetsky, F. Sun, M. Waterman, J. J. Chen, P. M. Chaudhary, J. Loscalzo, E. Crandall, and X. J. Zhou. 2014. ‘DiseaseConnect: a comprehensive web server for mechanism-based disease-disease connections’, Nucleic Acids Res, 42: W137–46.

Mann, D. M., A.M. McDonagh, J. Snowden, D. Neary, and S. M. Pickering-Brown. 2000. ‘Molecular classification of the dementias’, Lancet, 355: 626.

Mannino, D. M. 2002. ‘COPD: epidemiology, prevalence, morbidity and mortality, and disease heterogeneity’, Chest, 121: 121S–26S.

McClellan, J., and M. C. King. 2010. ‘Genetic heterogeneity in human disease’, Cell, 141: 210–7.

Menche, J., A. Sharma, M. Kitsak, S. D. Ghiassian, M. Vidal, J. Loscalzo, and A. L. Barabasi. 2015. ‘Disease networks. Uncovering disease-disease relationships through the incomplete interactome’, Science, 347: 1257601.

Mirnezami, R., J. Nicholson, and A. Darzi. 2012. ‘Preparing for precision medicine’, N Engl J Med, 366: 489–91.

Mistry, M., and P. Pavlidis. 2008. ‘Gene Ontology term overlap as a measure of gene functional similarity’, BMC Bioinformatics, 9: 327.

Murtagh, Fionn, and Pedro Contreras. 2012. ‘Algorithms for hierarchical clustering: an overview’, Wiley Interdisciplinary Reviews Data Mining & Knowledge Discovery, 2: 86–97.

Nakatsu, G., X. Li, H. Zhou, J. Sheng, S. H. Wong, W. K. Wu, S. C. Ng, H. Tsoi, Y. Dong, N. Zhang, Y. He, Q. Kang, L. Cao, K. Wang, J. Zhang, Q. Liang, J. Yu, and J. J. Sung. 2015. ‘Gut mucosal microbiome across stages of colorectal carcinogenesis’, Nat Commun, 6: 8727.

Newman, M. E. 2006. ‘Modularity and community structure in networks’, Proc Natl Acad Sci U S A, 103: 8577–82.

Okusaga, O., R. H. Yolken, P. Langenberg, M. Lapidus, T. A. Arling, F. B. Dickerson, D. A. Scrandis, E. Severance, J. A. Cabassa, T. Balis, and T. T. Postolache. 2011. ‘Association of seropositivity for influenza and coronaviruses with history of mood disorders and suicide attempts’, J Affect Disord, 130: 220–5.

Park, A. 2012. ‘Drugs zero in. Breast cancer, flu and obesity are in the crosshairs as drug companies produce more-targeted treatments’, Time, 179: 42.

Rzhetsky, A., D. Wajngurt, N. Park, and T. Zheng. 2007. ‘Probing genetic overlap among complex human phenotypes’, Proc Natl Acad Sci U S A, 104: 11694–9.

Serrano, M. A., M. Boguna, and A. Vespignani. 2009. ‘Extracting the multiscale backbone of complex weighted networks’, Proc Natl Acad Sci U S A, 106: 6483–8.

Sharma, A., J. Menche, C. C. Huang, T. Ort, X. Zhou, M. Kitsak, N. Sahni, D. Thibault, L. Voung, F. Guo, S. D. Ghiassian, N. Gulbahce, F. Baribaud, J. Tocker, R. Dobrin, E. Barnathan, H. Liu, R. A. Panettieri, Jr., K. G. Tantisira, W. Qiu, B. A. Raby, E. K. Silverman, M. Vidal, S. T. Weiss, and A. L. Barabasi. 2015. ‘A disease module in the interactome explains disease heterogeneity, drug response and captures novel pathways and genes in asthma’, Hum Mol, 24: 3005–20.

Sitas, F. 2016. ‘Twenty five years since the first prospective study by Forman et al. (1991) on Helicobacter pylori and stomach cancer risk’, Cancer Epidemiol, 41: 159–64.

Solovieff, N., C. Cotsapas, P. H. Lee, S. M. Purcell, and J. W. Smoller. 2013. ‘Pleiotropy in complex traits: challenges and strategies’, Nat Rev Genet Genet, 14: 483–95.

Tyner, C., G. P. Barber, J. Casper, H. Clawson, M. Diekhans, C. Eisenhart, C. M. Fischer, D. Gibson, J. N. Gonzalez, L. Guruvadoo, M. Haeussler, S. Heitner, A. S. Hinrichs, D. Karolchik, B. T. Lee, C. M. Lee, P. Nejad, B. J. Raney, K. R. Rosenbloom, M. L. Speir, C. Villarreal, J. Vivian, A. S. Zweig, D. Haussler, R. M. Kuhn, and W. J. Kent. 2017. ‘The UCSC Genome Browser database: 2017 update’ Nucleic Acids Res, 45: D626–D34.

Wang, K., H. Gaitsch, H. Poon, N. J. Cox, and A. Rzhetsky. 2017. ‘Classification of common human diseases derived from shared genetic and environmental determinants’, Nat Genet

Welter, D., J. MacArthur, J. Morales, T. Burdett, P. Hall, H. Junkins, A. Klemm, P. Flicek, T. Manolio, L. Hindorff, and H. Parkinson. 2014. ‘The NHGRI GWAS Catalog, a curated resource of SNP-trait associations’, Nucleic Acids Res, 42: D1001–6.

Wu, L., B. Zhou, N. Oshiro-Rapley, M. Li, J. A. Paulo, C. M. Webster, F. Mou, M. C. Kacergis, M. E. Talkowski, C. E. Carr, S. P. Gygi, B. Zheng, and A. A. Soukas. 2016. ‘An Ancient, Unified Mechanism for Metformin Growth Inhibition in C. elegans and Cancer’, cell, 167: 1705–18 e13.

Yang, J., and J. Leskovec. 2013. “Overlapping community detection at scale: a nonnegative matrix factorization approach.” In Proceedings of the sixth ACM international conference on Web search and data mining, 587–96. ACM.

Zanzoni, A., M. Soler-Lopez, and P. Aloy. 2009. ‘A network medicine approach to human disease’, FEBS Lett, 583: 1759–65.

Zhou, X., J. Menche, A. L. Barabasi, and A. Sharma. 2014. ‘Human symptoms-disease network’, Nat Commun, 5: 4212.

